# The response to single-gene duplication implicates translation as a key vulnerability in aneuploid yeast

**DOI:** 10.1101/2024.04.15.589582

**Authors:** H. Auguste Dutcher, James Hose, Hollis Howe, Julie Rojas, Audrey P. Gasch

## Abstract

Aneuploidy produces myriad consequences in health and disease, yet models of the deleterious effects of chromosome amplification are still widely debated. To distinguish the molecular determinants of aneuploidy stress, we measured the effects of duplicating individual genes in cells with varying chromosome duplications, in wild-type cells and cells sensitized to aneuploidy by deletion of RNA-binding protein Ssd1. We identified gene duplications that are nearly neutral in wild-type euploid cells but significantly deleterious in euploids lacking *SSD1* or *SSD1+* aneuploid cells with different chromosome duplications. Several of the most deleterious genes are linked to translation; in contrast, duplication of other translational regulators, including eI5Fa Hyp2, benefit *ssd1Δ* aneuploids over controls. Using modeling of aneuploid growth defects, we propose that the deleterious effects of aneuploidy emerge from an interaction between the cumulative burden of many amplified genes on a chromosome and a subset of duplicated genes that become toxic in that context. Our results suggest that the mechanism behind their toxicity is linked to a key vulnerability in translation in aneuploid cells. These findings provide a perspective on the dual impact of individual genes and overall genomic burden, offering new avenues for understanding aneuploidy and its cellular consequences.

## INTRODUCTION

Aneuploidy, the state of having an abnormal number of one or more chromosomes, is highly relevant to human health: it underlies embryonic inviability and trisomy syndromes, is associated with aging, and is a well-known feature of malignant cancers (1–3). Debates continue over the mechanisms driving the deleterious consequences of aneuploidy, specifically chromosome duplication of focus here. Models range from those concentrating on the increased generic burden of extra DNA and encoded protein (4, 5), to disruption of protein networks due to stoichiometric imbalance of dosage-sensitive proteins (6–8), to amplification of specific highly toxic gene products (9, 10). However, distinguishing between the deleterious effects of specific genes versus the collective burden of many genes is a significant experimental challenge, especially in mammalian systems. Thus, despite the relevance and prevalence of aneuploidy, we still lack a comprehensive understanding of the mechanisms behind the deleterious consequences of chromosome amplification.

Research in the yeast model organism, *Saccharomyces cerevisiae*, has expanded our understanding of the cellular impacts of aneuploidy and led to the discovery of key genetic vulnerabilities. Pioneering studies using the highly aneuploidy-sensitized laboratory strain W303 unveiled a range of defects in cells with extra chromosomes, including metabolic changes, cellular stress, defects in cell-cycle progression, and signs of protein aggregation and protein mismanagement known as proteostasis stress (4, 11–15). However, work from our lab and others has shown that the W303 is an outlier in its extreme sensitivity to chromosome duplication. In contrast to findings based on the W303 strain, aneuploidy is not uncommon in non-laboratory strains of yeast and found in ∼20% of strains distributed across the *S. cerevisiae* phylogeny. Strains with extra chromosomes typically grow substantially better than W303, though often with slower growth rates than isogenic euploid cells, in both wild yeast strains and a different lab strain derived from S288c (16–20). We previously showed that the W303 strain’s extreme sensitivity to chromosome duplication is due to a hypomorphic allele of the RNA-binding protein Ssd1 (21). Deletion of *SSD1* from wild strains recapitulates many of the aneuploidy phenotypes reported in W303, including signs of proteostasis stress (21). Ssd1 is thus a key mediator of aneuploidy tolerance, though its precise role in this modulation has remained elusive.

Ssd1, orthologous to human Dis3L2 but lacking nuclease activity, binds several hundred mRNAs via sequence motifs often found in the 5’ UTR (21–26). Bound mRNAs are enriched for a variety of encoded functions, including proteins localizing to the cell wall and proteins involved in cell cycle regulation, RNA metabolism, sterol transport, and a diverse collection of seemingly unrelated functions (21, 22, 24). Although molecular details are sparse, Ssd1 is reported to play a role in translation and mRNA localization to processing bodies and the cellular periphery, particularly for transcripts encoding cell wall proteins and other gene products localized to the nascent bud (22, 23, 26–31). Recent findings from our laboratory demonstrated that deleting *SSD1* sensitizes a wild strain to most chromosome duplications to varying degrees (32), indicating that the role of Ssd1 in aneuploidy tolerance is not limited to a specific chromosome. Thus, understanding the role of Ssd1 in tolerating chromosome amplification is a potent tool for investigating the molecular determinants of aneuploidy tolerance more broadly.

Separate from the role of Ssd1, the molecular determinants behind aneuploidy toxicity in functional cells are still widely debated. Work from *S. cerevisiae* transformed with yeast artificial chromosomes (YACs) containing human DNA inserts pointed to the importance of gene products—not merely the presence of extra DNA—as the cause of aneuploidy-related defects (4, 33). This fostered the idea that proteostasis stress in aneuploidy-sensitized strains is a driving determinant of aneuploidy toxicity (11, 34). One hypothesis is that simply having too much protein produces proteostasis stress. But another is that the over-abundance of specific proteins causes stoichiometric imbalance, namely proteins in multi-subunit complexes and those that participate in many protein-protein interactions (7, 35, 36). Both cases could cause a range of effects including protein aggregation (37), defects with protein quality control (11), and resulting osmotic challenges that disrupt endocytosis and other processes (38). At the other end of the spectrum, amplification of specific highly deleterious genes encoded on some chromosomes could pose particular problems. Given that whole chromosome amplification inherently increases the copy number of many genes simultaneously, distinguishing the effects of specific genes from generalized burden remains a complex task.

In this study, we leveraged Ssd1 to help distinguish between these possibilities. We used a yeast gene expression library to identify genes that are deleterious when duplicated in isolation, in euploid and aneuploid cells with and without *SSD1*. Assaying euploids and aneuploids in parallel enabled us to distinguish the baseline reactions to gene duplication in euploid cells from the unique outcomes when the same genes are amplified within an aneuploid context. This approach identified shared susceptibilities in Ssd1-deficient euploid cells and *SSD1+* aneuploids, allowing us to address disputed models of aneuploidy toxicity. We propose that defects associated with chromosome duplication are the result of an interaction between the generalized burden of chromosome amplification with specific genes whose duplication is largely neutral in euploid cells but become problematic in the aneuploid context. Our study demonstrates the value of leveraging genetic vulnerabilities to deepen our understanding of the intricate cellular state of aneuploidy and its extensive implications for human health.

## RESULTS

We set out to investigate the response of *ssd1Δ* cells to gene duplication. To do this, we used a collection of yeast strains recently generated in our lab, each harboring a duplication of a single *Saccharomyces cerevisiae* chromosome (32). This strain panel was engineered in a haploid derivative of oak-soil strain YPS1009 and includes all chromosome duplications except chromosome VI (Chr 6), which is known to be highly deleterious in other strain backgrounds (4, 39). In addition to aneuploid strains in the parental background, we derived a comparable panel in YPS1009 cells lacking *SSD1*; the only exception was duplication of Chr 16, which is unculturable in the absence of Ssd1. For most of the aneuploids tested, the fitness cost of the extra chromosome is significantly greater in the *ssd1Δ* background (32). Thus, Ssd1 contributes to aneuploidy tolerance across the majority of chromosome duplications, though to varying degrees by chromosome.

Chromosomes also varied in terms of their fitness cost when duplicated in wild-type cells. We wondered if the dependence on Ssd1 was proportionate to the inherent cost of each chromosome in the wild type. If so, it would suggest that Ssd1 is important for buffering, albeit incompletely, aneuploidy-induced stresses seen in the wild-type. An alternative model is that deletion of *SSD1* introduces entirely new sensitivities that are not relevant in the *SSD1+* aneuploids.

To distinguish these possibilities, we plotted the Ssd1 dependence of each chromosome duplication (scored as the relative growth rate of the *ssd1Δ* versus corresponding wild-type aneuploids) against the fitness defect of each chromosome duplication in the *SSD1+* wild-type (taken as the relative growth rate of each wild-type aneuploid relative to the euploid control, Fig 1). We found a modest yet significant correlation between Ssd1 dependence and chromosome fitness cost (R^2^ = 0.38, p = 8.3E-03). However, several chromosome amplifications were substantially more or less dependent on Ssd1 than predicted from the wild-type sensitivity. For example, Chr 12 duplication incurred one of the most severe defects in the absence of *SSD1*, surpassed only by Chr 16 duplication that is unculturable in the *ssd1Δ* strain. Conversely, Chr 2, Chr 14, and Chr 10 amplifications were proportionately less toxic than expected. Together these findings imply that Ssd1 has a generalizable role in coping with aneuploidy, but with some chromosome-specific effects. This indicates that Ssd1 deletion is a useful tool to exacerbate and interrogate determinants of aneuploidy toxicity.

**Figure 1.**
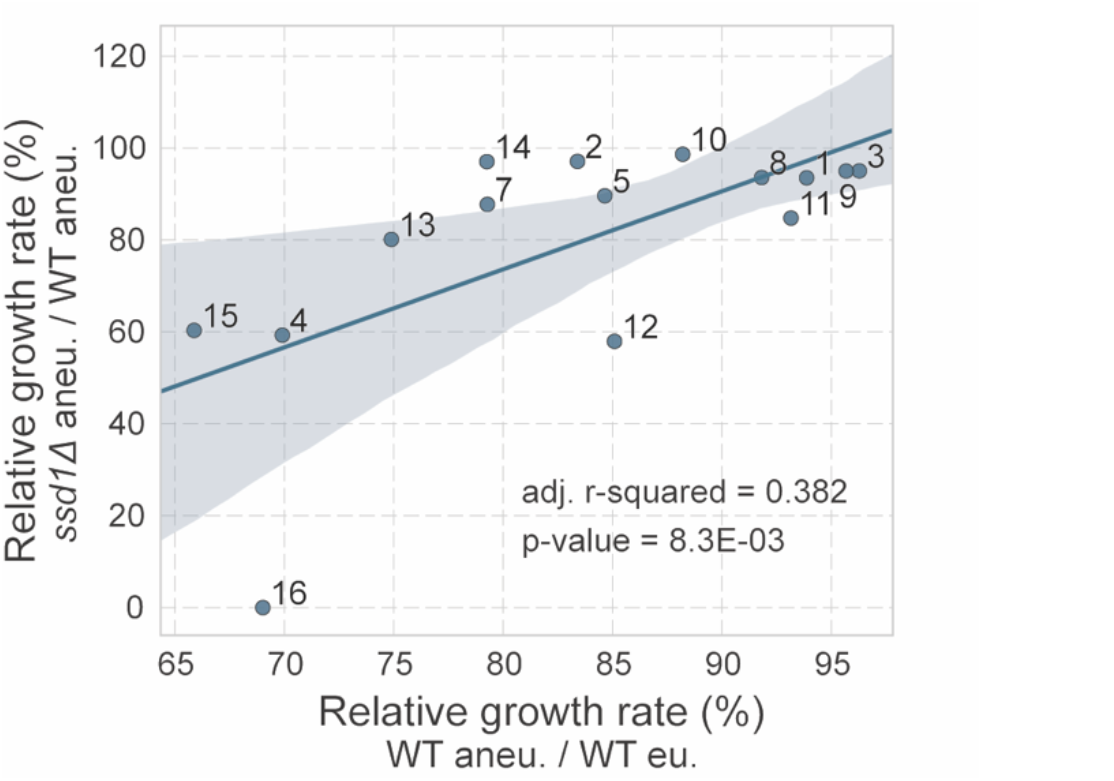
Ssd1 dependence is modestly correlated with aneuploidy toxicity in the *SSD1+* wild-type background. Each point represents the average relative growth rate of the *ssd1Δ* aneuploid versus wild-type aneuploid (y-axis) plotted against the average relative growth rate of wild-type aneuploid versus the euploid (x-axis), labeled to identify the amplified chromosome. Line shows ordinary least squares regression; shaded area denotes the 95% confidence interval (see Methods).

### Fitness costs of single-gene duplication measured in an aneuploid strain panel

Previous studies strongly suggested that aneuploidy toxicity is dependent on genes encoded on the amplified chromosomes (4, 33), but whether these effects, and Ssd1 dependence in particular, is dependent on only a few genes or many genes is unclear. To distinguish between models and study Ssd1’s role in aneuploidy tolerance, we used a plasmid library to measure the cost of duplicating genes individually. The MoBY 1.0 gene library comprises ∼5,000 open reading frames along with their native upstream and downstream sequences, cloned onto a barcoded centromeric (CEN) plasmid (40). We transformed the library into wild-type and *ssd1Δ* euploid strains as well as select wild-type and *ssd1Δ* aneuploids with duplications of Chr 4, Chr 7, Chr 12, and Chr 15 (the Chr 15 wild-type aneuploid was omitted due to poor library coverage). These strains were chosen because they present a range of growth defects and varying degrees of Ssd1 dependence. Duplicated chromosomes carried distinct selectable markers that were maintained in selective medium during competitive growth. Each pooled library was grown for 10 generations (with the exception of the YPS1009 Chr4 *ssd1Δ* strain, which was grown for only 5 generations due to its extreme fitness defect), in three or more biological replicates (see Methods). An aliquot of the library was collected before and after competitive growth, and the relative abundance of each plasmid was scored by sequencing the plasmid barcodes (see Methods). Barcodes that decrease in abundance during competitive growth correspond to genes whose duplication is deleterious, whereas barcodes that increase in abundance represent genes whose duplication is advantageous. To quantify fitness effects, we calculated a fitness score for each gene duplication, defined as the log_2_(fold change) of normalized barcode counts after competitive growth (see Methods).

We began by analyzing genes whose duplication affects the wild-type euploid strain. Based on seven replicates, we identified 2,403 genes whose duplication produced a significant fitness effect at a false discovery rate (FDR) < 0.05 ((32), see Methods and Supplemental Table 1). Nearly half of these duplications were beneficial (Fig 2A), and this group was enriched for essential genes (p = 9.1E-08, hypergeometric test). Genes whose duplication was detrimental were not enriched for any gene ontology terms, but the group was enriched for genes encoding longer proteins and proteins with a higher proportion of disordered regions (32), consistent with previous reports (41). There was no enrichment for proteins in multi-subunit complexes and no difference in the number of protein interactions for deleterious genes compared to neutral genes. None of the chromosomes showed a significant overrepresentation of detrimental genes (FDR > 0.05).

**Figure 2.**
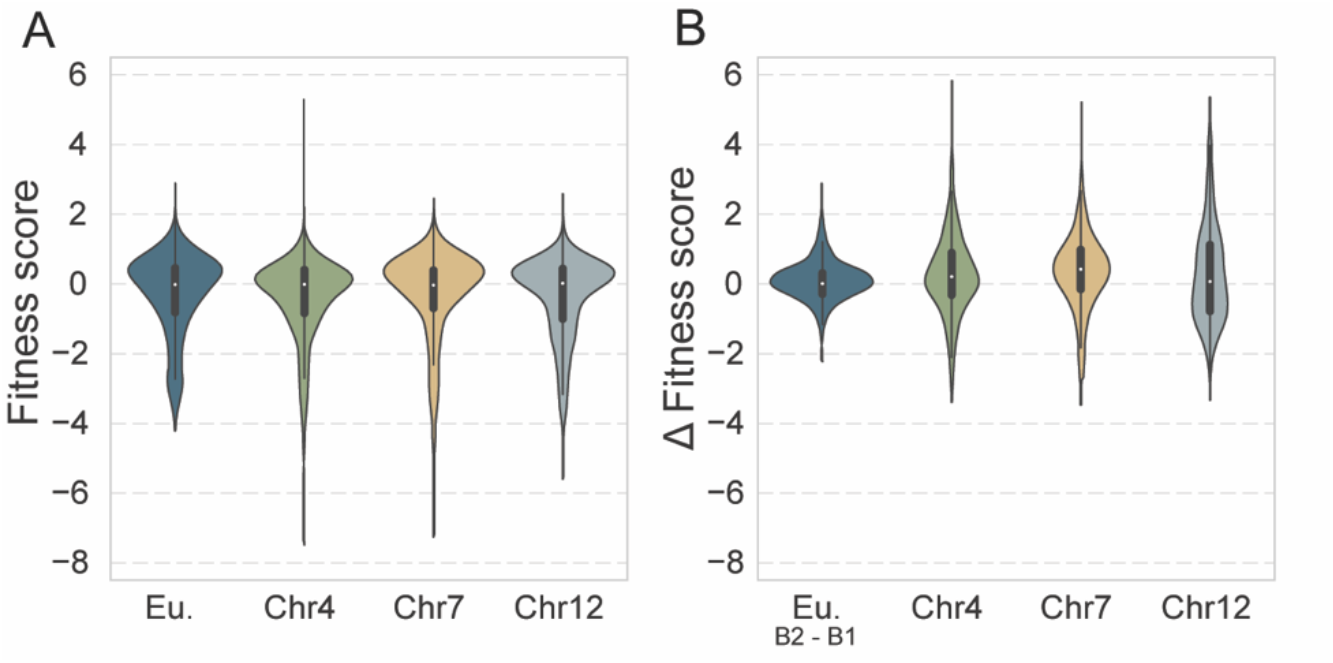
Distribution of fitness costs of gene duplication are similar in aneuploids and euploids. A) Distribution of fitness scores (log_2_(fold change) of normalized barcode counts) for all measured genes in euploid (Eu.) and aneuploids with the denoted chromosome amplification. B) Distribution of pairwise differences in fitness scores between each wild-type aneuploid and batch-paired euploids, for 2,403 detrimental genes identified in the euploid wild-type. As a control, the euploid distribution represents the difference in fitness scores between two batches (B1 and B2) of euploid replicates.

### Aneuploid strains are not universally more sensitive to gene duplications

With the wild-type euploid response to gene duplication as a baseline, we next assessed the effects in aneuploid strains. At the outset of the experiment we hypothesized that aneuploids may be more sensitive to the gene duplication library, since they already carry a burden of extra gene content. If aneuploids were more sensitive to gene duplication, we would expect more deleterious genes in aneuploid strains and/or more severe fitness defects upon duplication. On a global scale, this was not the case: the proportion of detrimental genes was comparable between aneuploids and batch-paired euploids, albeit with a marginally higher proportion of detrimental genes in aneuploids (Fig 2A). This could not be explained by data normalization issues, since we confirmed that genes scored as neutral in the library experiment had no significant effect on growth rate when strains were grown in isolation (Supplemental Fig 1). Although there was a longer tail of deleterious genes in the aneuploid strains, the bulk of the distribution was similar to euploids, and the majority (87-90%) of fitness scores in aneuploids were within 1.5 fold of batch-paired euploids. We conclude that the aneuploid strains are not generally sensitized to further gene over-production but do exhibit increased sensitivity to a subset of duplicated genes.

To investigate further, we identified 335 genes that were deleterious in one or more aneuploids (FDR < 0.05) and at least 1.5 fold more deleterious than batch-paired euploid measurements (see Methods). Of these 335 genes, 58 were common to two or more aneuploid strains, suggesting common effects across chromosomes. There was no enrichment for any gene ontology terms. However, we noticed that 8 of the common detrimental genes are naturally encoded adjacent to a centromere, such that the centromere was cloned onto the plasmid along with the gene. Plasmids with a cloned centromere are overrepresented among genes more deleterious in aneuploids (p=5.5E-09, hypergeometric test), raising the possibility that it is the cloned centromere that is detrimental and suggesting that aneuploids may have a defect managing plasmids with multiple centromeres. In contrast, we noticed that some genes scored as deleterious in the euploid were much less deleterious in aneuploid strains (see Fig. 2B).

We identified 360 genes that were deleterious in the euploid (FDR < 0.05) but at least 1.5 fold less so in one or more wild-type aneuploids. 153 of these genes were common to two or more aneuploid strains, again suggesting common effects independent of the specific chromosome duplicated. Intriguingly, these common genes were enriched for genes involved in the mitotic spindle (p = 9.1E-05). Genetic perturbation of mitotic spindle genes was previously shown to be deleterious in polyploid yeast (42), thus our results hint that aneuploids may benefit from increased abundance of spindle components. In sum, our results suggest that aneuploid strains are not generally more sensitive to further gene duplication but that a subset of genes are more or less toxic in aneuploids.

### Specific gene duplications are especially toxic in ssd1Δ euploids and explain a portion of Ssd1 aneuploidy dependence

As shown above, *SSD1* deletion sensitizes cells to aneuploidy to varying degrees depending on which chromosome is amplified. Given Ssd1’s role in translational suppression, one question was if *SSD1* deletion sensitizes cells to gene over-production in general. We therefore measured fitness costs of gene duplication in the *ssd1Δ* euploid. Overall, the distribution of fitness scores for gene duplications was not significantly different than the euploid wild-type (p = 0.38, Wilcoxon rank-sum test, Fig 3A), indicating that *ssd1Δ* does not sensitize cells to all gene duplications. However, we did identify 218 genes that were significantly more detrimental in *ssd1Δ* euploid versus wild-type euploid strains (FDR < 0.05, Fig 3B, Supplemental Table 3). We were especially interested in identifying functional or biophysical properties of the encoded products; however, the group did not have striking functional enrichments, aside of 4 genes involved in ER to Golgi transport (p = 1.7E-03, hypergeometric test), nor differences in more than a dozen other transcript or protein features including abundance, structure or sequence composition, or other biophysical properties (see Methods). Furthermore, the group was not enriched for mRNAs bound by Ssd1 (p = 0.67, hypergeometric test) or that harbor the known Ssd1 binding motif (22, 24) or any other sequence motif. Concordantly, the group of genes encoding Ssd1-bound mRNAs was not more deleterious in the *ssd1Δ* euploid strain (Fig 3C). The only significant feature among the 218 genes was a weak enrichment for genes that are shorter than all other genes in the dataset (p = 1.3E-04, Mann-Whitney U). This was particularly notable since genes whose duplication is detrimental in the euploid wild-type strain are enriched for longer genes (discussed in (32)). Without a clear signature from any of the tested features, the reason behind the toxicity of these genes in *ssd1Δ* cells is unknown. Nonetheless, some genes are reproducibly more deleterious when duplicated in an *ssd1Δ* context.

**Fig 3.**
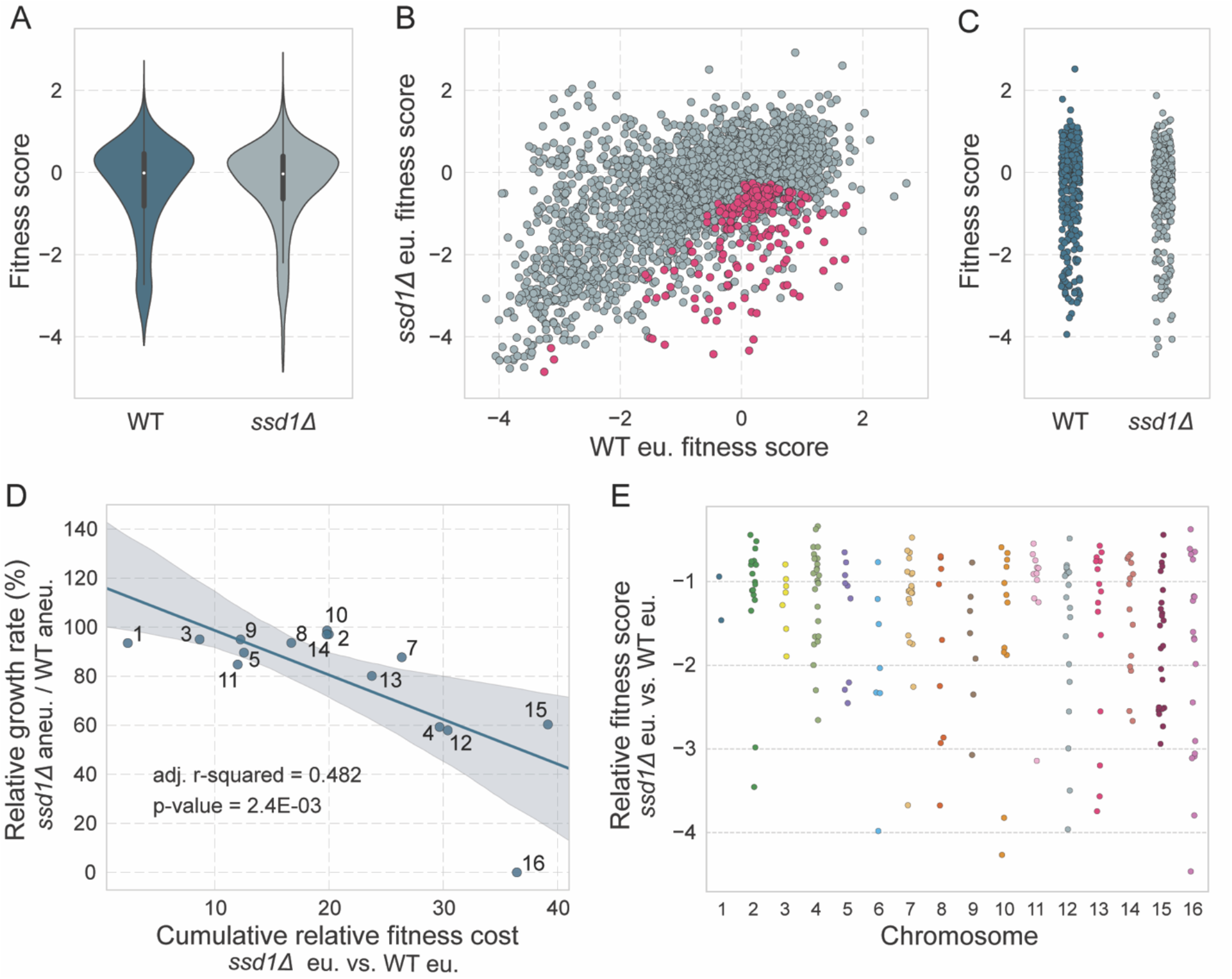
A subset of genes are more detrimental in *ssd1Δ* euploid cells relative to wild type. A) Distributions of fitness scores for all 4,462 measured genes in wild-type euploid and *ssd1Δ* euploid strains. B) Scatterplot of replicate-averaged fitness scores for all measured genes in *ssd1Δ* euploid (y-axis) and wild-type euploid (x-axis); 218 genes significantly more deleterious in *ssd1Δ* euploid cells are shown in pink. C) Fitness scores for 336 genes encoding mRNAs bound by Ssd1 (see Methods) in wild-type (blue) and *ssd1Δ* (gray) euploid strains. D) Average relative growth rate of the *ssd1Δ* aneuploid versus wild-type aneuploid (y-axis, as shown in Fig 1) plotted against the sum of replicate-averaged relative fitness costs (*ssd1Δ* euploid versus *SSD1+* euploid) for the subset of 218 genes whose duplication is toxic to *ssd1Δ* cells (x-axis). E) Ssd1-dependent fitness scores for the 218 genes represented in (D), plotted according to encoding chromosome.

We next set out to model Ssd1 dependence using the fitness costs measured for these individual gene duplicates. In other recent work from our lab, we showed that much (up to 69% of the variance) of the fitness costs of chromosome duplication in a wild-type strain can be explained by the additive fitness costs of all genes duplicated on each chromosome (32). Here we focused on explaining the degree to which each chromosome duplication depends on Ssd1 for tolerance. We calculated the per-chromosome Ssd1 dependence as the sum of per-gene Ssd1 dependences, *i*.*e*. the sum of relative fitness costs of each gene duplicated in the *ssd1Δ* euploid versus in the wild-type euploid strain, considering the 218 genes identified above whose duplication is especially deleterious in the *ssd1Δ* euploid strain. This simple model explained roughly half of the variance in Ssd1 aneuploidy dependence, with an adjusted R^2^ of 0.48 (Fig 3D, p-value: 2.4E-03). Omitting Chr 16 from the analysis increased the explanatory power to 52% (adj. R^2^ 0.52). Model performance decreased substantially when only the top 10th percentile of *ssd1Δ*-sensitive genes were used (36% of variance explained, Supplemental Fig 1). Thus, the cumulative burden of specific genes on each chromosome explains roughly half of the variance in Ssd1 dependence, but this set is more than a small handful of genes.

### Many gene duplicates toxic in ssd1Δ euploids are also toxic to SSD1+ aneuploids but not SSD1+ euploids

We hypothesize that Ssd1 is required to buffer the stress of chromosome duplication; thus, we reasoned that *SSD1+* aneuploids may show sensitivity to the same gene duplications that are problematic in the *ssd1Δ* euploid strain. We explored this in several ways. First, on further investigation we realized that the group of genes more deleterious in one or more aneuploid strains is enriched for the 218 genes whose duplication is especially deleterious in *ssd1Δ* euploid cells (p = 2.9e-06, hypergeometric test). Second, we plotted for these 218 genes the aneuploid-averaged fitness scores from the three wild type aneuploids against the euploid fitness scores (Fig 4A). Indeed, more of the genes’ scores were deleterious in aneuploids than in the euploid (145 genes with negative values in aneuploids versus 69 genes in the euploid). One subset of genes was especially toxic in aneuploids compared to the euploid strain, indicated by their deviation from the linear trend (Fig 4A, labeled points). We confirmed that these duplicated genes showed significantly higher fitness costs in most of the individual aneuploids compared to the euploid (Fig 4B). We conclude that many of the gene duplications toxic in *ssd1Δ* euploids are also more deleterious in *SSD1+* aneuploids versus the euploid (see Discussion).

**Figure 4.**
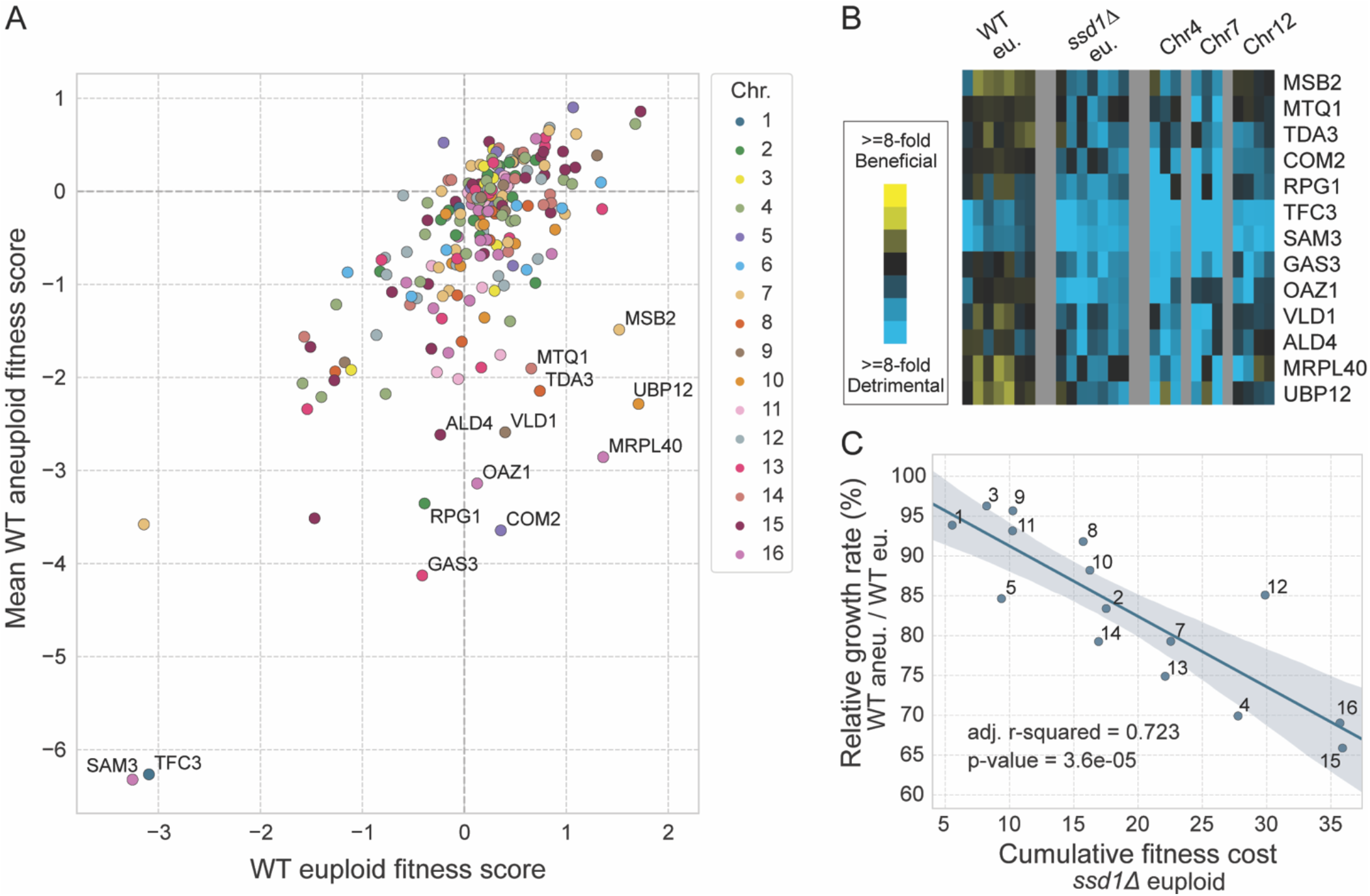
Many gene duplications toxic in *ssd1Δ* euploids are also toxic in *SSD1+* aneuploids. A) Mean aneuploid fitness scores for each gene scored across wild-type (WT) aneuploids (y-axis) plotted against the corresponding fitness scores measured in wild type euploid cells (x-axis), for the 218 genes in Fig 3B. Genes with especially deleterious effects in aneuploids relative to the euploid are labeled with gene name. B) Fitness scores for genes labeled in Fig 4A (rows) in biological replicates of each strain (columns), colored according to the key. C) Average relative growth rates of wild-type aneuploids versus euploid (y-axis) plotted against the sum of log_2_ fitness costs for the 218 gene duplicates as measured in the *ssd1Δ* euploid strain (x-axis).

We wondered if the additive cost of duplicating these 218 genes could explain *SSD1+* wild-type aneuploid fitness defects. An additive model based on individual-gene fitness costs measured in the euploid wild-type strain was a poor predictor of the measured costs of chromosome duplication in the *SSD1+* background (adjusted R^2^ = 0.06, p-value = 0.19). But many of these genes are only deleterious when duplicated in the *ssd1Δ* euploid strain. We therefore modeled the fitness cost of chromosome duplication in *SSD1+* cells but based on the additive cost of gene duplications measured in the euploid *lacking SSD1*—remarkably, this model explained 72% of the variance in growth rates of the *SSD1+* aneuploids (Fig 4C). This important result indicates that chromosome duplication in an *SSD1+* background likely mimics the stress of *SSD1* deletion in a euploid strain, sensitizing cells to the same subset of problematic gene duplications, likely through the same underlying mechanisms (see Discussion).

### Duplicated genes beneficial in ssd1Δ aneuploids implicate mechanisms of aneuploidy tolerance

Since *SSD1* deletion sensitizes strains to the stress of chromosome duplication, we wondered if genes beneficial in *ssd1Δ* aneuploids might complement processes that are defective in those aneuploids. We therefore identified genes whose duplication compensates growth defects in *ssd1Δ* aneuploids (Fig 5A, see Methods). We identified the greatest number of beneficial genes in *ssd1Δ* cells with an extra copy of Chr12, possibly due to increased statistical power from 4 replicates, followed by *ssd1Δ* Chr4 aneuploid cells, and the fewest beneficial duplications in the *ssd1Δ* Chr7 aneuploid, the least dependent on Ssd1 of the aneuploids studied here. We identified 445 genes whose duplication was beneficial to at least one strain (FDR < 0.05). To validate, we tested 13 plasmids expressed in Chr12 aneuploids; 8 of the 13 genes significantly improved growth in the *ssd1Δ* aneuploid relative to empty vector, confirming library measurements (Supplemental Fig 1).

**Figure 5.**
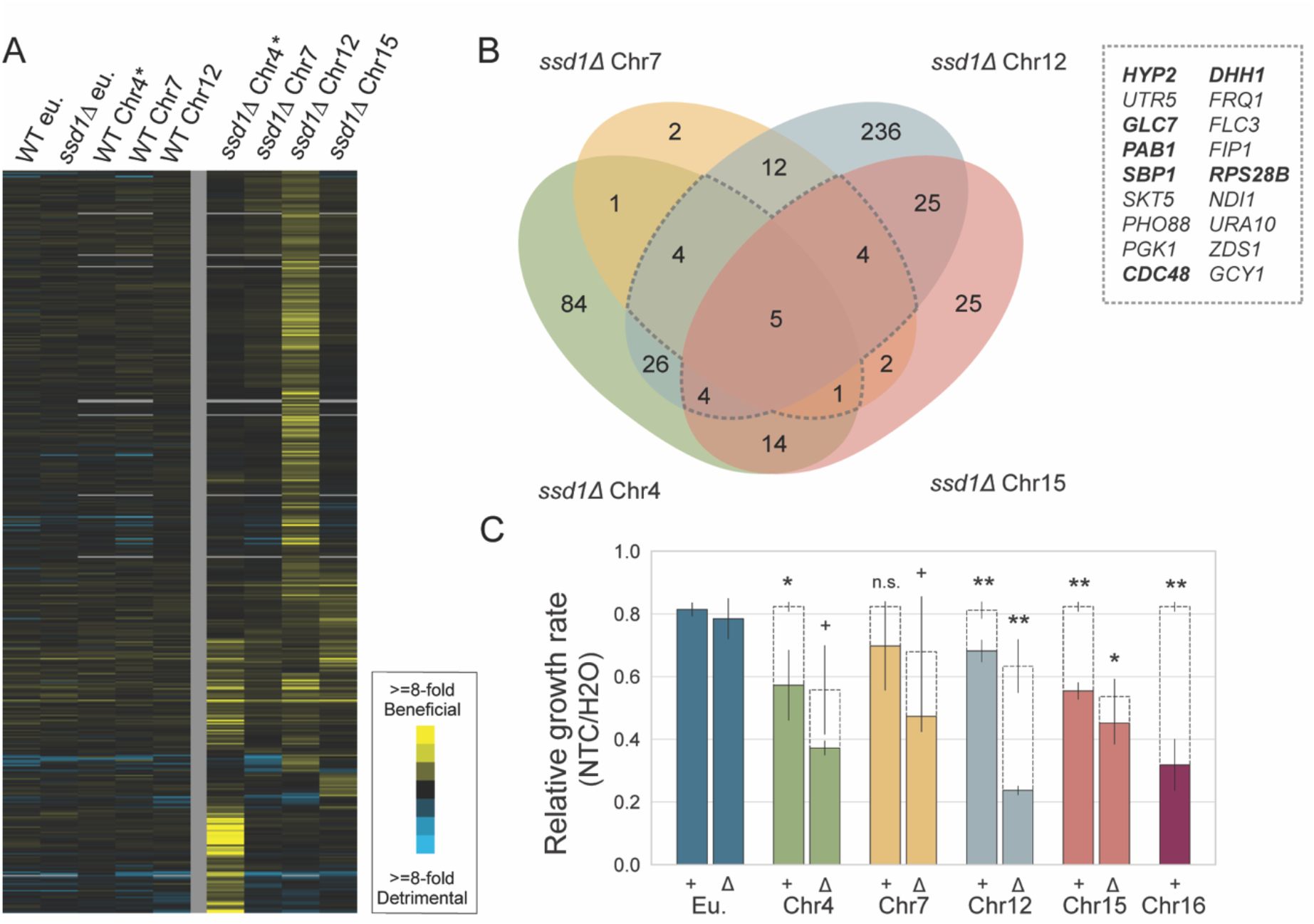
Gene duplications that benefit *ssd1Δ* aneuploids implicate translation. A) Hierarchically clustered fitness scores for 445 gene duplications (rows) beneficial in at least one aneuploid strain (columns), colored according to the key. Fitness scores represent the average of at least biological triplicate. (*) denotes scores measured after 5 generations instead of 10 generations, see text. B) Venn diagram showing the overlap of beneficial genes identified in each *ssd1Δ* aneuploid. Dotted border identifies beneficial genes common to 3 or more *ssd1Δ* aneuploids; bolded genes have known functions linked to translation. C) Average and standard deviation of relative growth rates of the denoted strains grown in 1ug/mL nourseothricin (NTC) compared to a water control (n=3). Measured growth rates are indicated as solid bars; expected growth rates based on the combined effect of NTC and underlying aneuploidy growth defect are shown as dashed bars (see Methods). (**) p-value < 0.01, (*) p < 0.05, (+) p < 0.1, one-tailed paired T-test.

Comparing across strains revealed that many gene duplications were beneficial to multiple *ssd1Δ* sensitized aneuploids (Fig 5B, see Methods). We were especially interested in these genes, which may point to processes that are generally compromised upon chromosome duplication. 98 of these gene duplications showed positive fitness scores in at least 2 aneuploids, and 18 duplications were positive in at least 3 aneuploids. The group of 98 was enriched for genes encoding small GTPase-mediated signaling proteins (p = 3.7E-05, hypergeometric test) and mRNA processing proteins (p = 9.8E-04), while the group of 18 was enriched for translational regulators (p = 1.1E-03, hypergeometric test). Somewhat surprisingly, we did not find enrichment of genes linked to protein folding or the ubiquitin-proteasome system, which were previously associated with aneuploidy tolerance in the *ssd1-* W303 strain (43). Of the five plasmids with positive fitness scores in all four aneuploid *ssd1Δ* strains, four encode proteins directly implicated in translational regulation, including poly-A binding protein Pab1, translational repressor Sbp1, and eIF5a elongation factor Hyp2 (which was also cloned on a plasmid nominally carrying the adjacent gene, *UTR5*). The fifth gene, *GLC7*, encodes PP1 phosphatase that can act on translational regulator eIF2a in yeast (44), as well as other proteins. We confirmed that *PAB1, SBP1*, and *HYP2* all significantly improve growth rates of *ssd1Δ* Chr12 aneuploid cells relative to empty vector (p < 0.05, Suppl. Fig 3A). Additionally, *HYP2* duplication improved growth in all 4 *ssd1Δ* aneuploids relative to their corresponding wild-type aneuploids (Supplemental Fig 3B). These results strongly suggest that *ssd1Δ* aneuploid strains share defects that can be complemented by over-expressing specific genes involved in translation.

In retrospect, we realized that several of the genes whose duplication was especially *deleterious* in aneuploid cells are also connected to translation and Hyp2 specifically (Fig 4). Among those genes was *OAZ1*, which encodes an anti-enzyme that negatively regulates polyamine synthesis (45). Polyamines are precursors of hypusine, a non-canonical amino acid required for Hyp2 function (Schnier et al. 1991; reviewed in Park and Wolff 2018). Oaz1 production is regulated via a programmed ribosomal frameshift as well as a programmed ribosome stall (45, 48), which together serve as readouts of translational fidelity that is in part dependent on Hyp2 (49). Overproduction of Oaz1 is expected to indirectly inhibit Hyp2 function by downregulating the synthesis of polyamines required for hypusination. Intriguingly, *OAZ1* is encoded on Chr16, whose duplication is essentially inviable in the *ssd1Δ* background. Several other genes from Fig 5B are also linked to these processes: *SAM3* involved in S-adenosylmethionine (SAM) and polyamine transport (50), *MTQ1*, a SAM-dependent translational regulator (51), and *RPG1*, a subunit of eIF3 with roles in translation reinitiation as well as stop codon readthrough (52, 53). Together, these links reinforce the idea that *ssd1Δ* aneuploid strains and, to a lesser extent wild-type aneuploids, may have defects in translation.

Our previous work showed that the Chr12 aneuploid and especially *ssd1Δ* Chr12 aneuploid are sensitive to nourseothricin (NTC) (21), a translation-elongation inhibitor that at low doses stalls ribosomes, induces tRNA misincorporation and produces protein misfolding (54, 55). To test if this is a generalizable feature of aneuploid strains, we investigated other chromosome duplications for sensitivity to NTC. Indeed, all *SSD1+* aneuploids grew significantly slower than expected based on an additive model of aneuploidy and NTC effects (p < 0.05 by paired t-test, Fig 6C, see Methods) except the Chr7 aneuploid, which missed the significance cutoff. On the other end of the spectrum, duplication of the highly Ssd1-dependent Chr16 produced the greatest sensitivity to the drug. All of the *ssd1Δ* aneuploids treated with NTC grew reproducibly slower than expected across all replicates. In summary, these aneuploid strains showed above-additive sensitivity to translation elongation inhibitor NTC. Together with the results from our gene duplication screen, this sensitivity implicates translation as a particular vulnerability of wild-type aneuploids, which is exacerbated in the absence of *SSD1*.

## DISCUSSION

Although a rich body of literature describes phenotypes associated with chromosome duplication, the molecular causes of those phenotypes and aneuploidy toxicity in general remain poorly understood. Previous studies suggest that the products of amplified genes are to blame (4), but whether this is due to specific products and their associated properties or functions, or to a general burden associated with the increased genic load remains unknown. Many studies of Down syndrome (DS) focus on single genes amplified on human chromosome 21 (9, 10), and the effects of these genes are often studied via duplication in isolation in euploid tissue culture. On the other hand, several lines of evidence instead implicate the generalized burden of aneuploidy, independent of which chromosome is amplified. For example, Bonney *et al*. found that extra copies of a few dosage-sensitive genes expressed on each chromosome were not sufficient to explain reduced proliferation of aneuploids in a sensitized laboratory strain (5). Several other studies emphasize consequences of aneuploidy that are karyotype-independent (4, 11, 13, 14, 38, 56–58). Many perspectives suggest that it is some combination of the two models, where deleterious genes on each chromosome combine with increased sensitivity of aneuploid strains that have an increased underlying burden on cellular systems.

Our results provide evidence for an important and distinct model: that the generalized burden of chromosome duplication sensitizes cells to the duplication of specific genes on those chromosomes that are not otherwise deleterious when duplicated on their own but become so in the aneuploid context. This model posits a genetic interaction between generalized and gene-specific effects of chromosome duplication in *SSD1+* cells. We propose that the generalized effect of chromosome amplification mimics to some degree the vulnerability of *ssd1Δ* euploid cells to specific gene duplications, and that the physiological limitations of *ssd1Δ* cells, namely those related to translation (perhaps among other things), are also at play in *SSD1+* aneuploids.

Several lines of evidence support this model. On the one hand, we found that the growth defects incurred by chromosome duplication in both wild-type and *ssd1Δ* cells can be party explained by a subset of toxic genes. Importantly, this group does not simply represent the most problematic genes in the wild-type euploid; in fact, many are near-neutral when duplicated in the euploid wild type. Instead, the cellular context created by *SSD1* deletion or chromosome amplification in *SSD1*+ cells induces a specific vulnerability that renders these duplications toxic. In strains with chromosome amplifications, this vulnerability could itself be created by a small subset of specific genes amplified on the chromosomes; however, given the increased sensitivity to these gene duplications across multiple chromosome duplications, we favor the hypothesis that it is the generalized burden of aneuploidy arising from the cumulative effect of amplifying many genes at once (32). The model presented here—that particular genes become problematic in the context of aneuploidy—has important implications for studies on aneuploidy toxicity, including the etiology of human aneuploidy syndromes. For example, many DS studies interrogating single genes on human chromosome 21 probe effects through single-gene duplication in euploid cell lines (59–63). Our results raise caution about modeling these phenotypes in euploids and, as other have noted (64), underscore the importance of considering the cellular context of aneuploidy.

Despite these important insights, several unanswered questions remain. Why do certain genes become deleterious in this specific context and what is the precise role of Ssd1 in their detrimental effects? Among the 218 gene duplications especially deleterious in *ssd1Δ* euploid cells, the only shared signature we identified among them is that many are shorter than other genes. Short transcripts associate more with “closed loop” factors involved in mRNA circularization, which facilitates faster translation reinitiation (65, 66). This could be related to translational effects discussed below. Notably, we did not find these Ssd1-dependent genes to be more highly expressed at the RNA or protein level, nor did we find overrepresentation of genes encoding protein complexes, proteins with many protein-protein interactions, or proteins with a high propensity for disorder. Work from our lab and others (32, 67) strongly suggest that genes encoding multi-subunit complexes are not toxic when merely duplicated, likely because cells have evolved mechanisms to manage their moderate increase (16, 32, 68, 69). Therefore, the harmful effects identified in this study likely arise from unrecognized attributes, necessitating more research to investigate further.

Additionally, the mechanism by which Ssd1 influences these genes’ toxicity, if not directly regulating their mRNAs, remains unclear. Ssd1 interacts physically with the ribosome and other proteins that regulate translation (including Pab1 identified above as beneficial to *ssd1Δ* aneuploids) (70, 71) and it directly binds hundreds of mRNAs through direct contact (24). One possibility is that Ssd1 has a general role in translation at many more mRNAs than those bound through direct contact. But another possibility is that misregulation of transcripts bound directly by Ssd1 could trigger secondary widespread effects on translation dynamics or fidelity, resulting in toxicity effects at duplicated non-target genes. Examples of such widespread effects exist in the literature. For instance, Aviner et al. (72) demonstrated that the ribosome collisions on mutant Huntington transcript produce a cycle of dysfunction that sequesters eIF5a, thereby depleting it from other transcripts and altering translation of stress-responsive transcripts, in part by misregulating their upstream open reading frames (uORFs). Interestingly, a recent preprint linked yeast gene copy number variation, Ssd1, and uORFs by suggesting that genes whose encoded mRNA abundance is substantially discordant with its protein abundance were more likely to contain uORFs, and that those reading frames were enriched for Ssd1 binding motifs (73). While we were unable to find enrichment of known uORFs among the 218 genes, these findings highlight how the altered regulation of Ssd1 targets could render a specific group of gene duplicates toxic, through direct or indirect effects.

Other lines of evidence support that aneuploidy creates a context in which translational efficiency and/or fidelity is perturbed. Both the sensitized lab strain and wild strains lacking *SSD1* are highly sensitive to translation inhibitors including cycloheximide, G418 and/or NTC, but only in the context of extra chromosomes (4, 21, 43). This is consistent with a role for Ssd1 in translation and the idea that chromosome amplification produces translational stress that requires Ssd1 to manage (26, 30). But wild strains with functional *SSD1* are also more sensitive to NTC than euploid strains (21), suggesting that this vulnerability, while significantly exacerbated in the absence of Ssd1, exists also in *SSD1+* cells. Studies in other organisms show that translational efficiency is also perturbed in response to monosomy, apparently driven by imbalanced expression of ribosomal proteins (74, 75); thus karyotype imbalance more broadly could also perturb global translation. Also supporting a key vulnerability in protein synthesis, we found that duplication of genes involved in translation, notably eIF5a (*HYP2*), ameliorates the growth defect of *ssd1Δ* aneuploids relative to corresponding wild-type aneuploids. In addition to its roles in elongation (76–79), eIF5a plays a key role in preventing ribosome stalling at stretches of poly-proline and other residues as well as at stop codons (80–82). Interestingly, yeast proteins particularly dependent on eIF5a include several GTPases (82), a group enriched among commonly beneficial genes identified here. We also found links to eIF5a among genes detrimental to the growth of *ssd1Δ* euploid and aneuploid cells. For example, full-length Oaz1 represses polyamine synthesis, and thus duplication of the *OAZ1* gene, especially if translational control that limits its abundance is disrupted (49), could limit polyamine availability for essential Hyp2 hypusination, which could in turn exacerbate global translation elongation stress. Intriguingly, SAM metabolism was recently found to be defective in *Drosophila melanogaster* lines with ploidy changes (83); thus, eIF5a-related pathways identified here may have relevance for understanding aneuploidy tolerance across systems.

An intriguing idea is that aneuploidy-induced translational stress provides the unrecognized source of proteostasis stress in aneuploid cells. Once again, Ssd1 may aid us in generating hypotheses. Strains lacking fully functional Ssd1 are more vulnerable to defects in the Elongator complex, which is essential for chemically modifying specific tRNA (29). Lack of these modifications results in less efficient selection of the correct tRNA, tRNA misincorporation, frameshifting, and increased protein aggregation (84–87). This is consistent with the increased sensitivity of *ssd1Δ* aneuploids to aminoglycosides like NTC, which directly bind the ribosome to cause tRNA misincorporation, and subsequently protein misfolding and aggregation (54). Indeed, we previously showed that NTC treatment of *SSD1+* aneuploids mimics signatures of protein aggregation found in *ssd1Δ* aneuploids (21). While having too many proteins can burden the cellular homeostasis network, it is likely that proteostasis management becomes even more challenging in the face of translational errors and inefficiencies that may be created by aneuploidy. This may compound to further overwhelm protein folding and maintenance systems leading to compromised proteostasis. In this context, Ssd1 emerges as a critical factor to manage translational stress. This link between disruptions in elongation and protein aggregation underscores how changes in the dynamics of translation can profoundly affect aneuploid cell physiology. Investigation of precisely how translational regulation intersects with proteostasis in aneuploids represents a fruitful area of ongoing inquiry.

## METHODS

### Strains and growth conditions

Further information about aneuploid panel, including strains used for growth phenotypes under standard conditions, can be found in (32). All other strains used in this study are listed in Supplemental File 1. Strains used in gene duplication experiments were transformed with a pool of the molecular barcoded yeast ORF library (MoBY 1.0) containing 5,037 barcoded CEN plasmids (40). At least 25,000 transformants were scraped from agar plates for roughly fivefold replication of the library, and frozen glycerol stocks were made. Multiple independent transformations of the pooled library were performed for each strain (see competitive growth details below). Competitive growth was done in liquid synthetic media lacking histidine (SC-His) and with NTC (Werner BioAgents and Jena Bioscience, 100 mg/L) to maintain aneuploidy, with G418 (Research Products International, 200 mg/L) added for plasmid selection. Experiments interrogating single genes were performed using culture growths in test tubes at 30°C with shaking. For NTC experiments, cells were cultured 3 hours into log phase; 1ug/mL NTC or an equivalent volume of water (vehicle control) were added; OD_600_ measurements recorded beginning 1 hour after NTC addition. Expected growth rates were calculated using batch-paired growth rates as follows: the expected growth rate (GR) of each aneuploid treated with NTC was taken as the % change in euploid growth rate after NTC treatment; the expected GR of each *ssd1Δ* aneuploid treated with NTC was taken as the product of the % reduction in growth of the corresponding wild-type aneuploid treated with NTC and the % reduction in growth of each *ssd1Δ* aneuploid versus corresponding wild-type aneuploid in the absence of NTC treatment.

### MoBY 1.0 competitive growth

Competition experiments were performed as in (40, 88). Briefly, 1 mL frozen glycerol stocks of library transformed cells were thawed into 100 mL of liquid SC-His with NTC (100 mg/L) and G418 (200 mg/L) at a starting OD_600_ of 0.05, then grown in shake flasks at 30°C with shaking. An aliquot of frozen stock represented the starting pool (generation 0) for each strain. After five generations, each pooled culture was diluted to an OD_600_ of 0.05 in fresh media to maintain cells in log phase. At 10 generations (5 generations where indicated) cells were harvested and cell pellets were stored at −80°C.

Data were collected in two batches. Batch 1 data was generated from two independent competitive growths each from two independent transformations of the plasmid library, for a total of 4 replicates. Reproducibility was consistently high between independent growths from the same transformation stock (Pearson correlation coefficients PCC 0.87-0.96) and more modest across independent growths from different transformation stocks (PCC 0.44-0.67). Thus, to capture this variation, each of the three independent growths in Batch 2 was performed from an independent transformation of the pooled library. This ultimately yielded noisier fitness scores but greater confidence in reproducible effects. We pooled euploid controls from Batch 1 and Batch 2 for 7 replicates each of the wild-type euploid and *ssd1Δ* euploid experiments. Fitness scores generated from Batch 1 and Batch 2 were reproducible, with PCC of 0.93 and 0.92 for wild-type euploid and *ssd1Δ* euploid, respectively.

### MoBY 1.0 sequencing

Plasmids were recovered from each pool using Zymoprep Yeast Plasmid Miniprep II (Zymo Research D2004-A). Plasmid barcodes were amplified using primers described in (89) and 20 PCR cycles. Barcode amplicons were pooled and purified using AxyPrep Mag beads (1.8X volume beads per sample volume) according to the manufacturer’s instructions (Axygen). Pooled amplicons were sequenced on one lane of an Illumina HiSeq 4000 to generate single-end 50 bp reads. This resulted in 126.2 million reads with exact matches to MoBY 1.0 barcodes in Batch 1 across 32 samples (average of ∼3.9M reads/sample) and 154.7 million reads in Batch 2 across 51 samples (average of ∼3.0M reads/sample). Sequencing data and barcode counts are available in the Gene Expression Omnibus (GEO) under accession GSE263221.

### MoBY 1.0 dataset analysis

To limit bias from barcodes that were under-represented in the starting pool, we removed barcodes in the bottom 5% of total counts at generation zero or with 0 reads at generation 0 in any sample. A pseudocount of 1 was added to every measurement. Data were normalized by TMM normalization (90) in edgeR version 3.36.0 (91), which produced very similar results to normalization by total barcodes sequenced. Barcodes that changed significantly in abundance in each experiment were identified using a genewise negative binomial generalized linear model with quasi-likelihood tests, taking FDR < 0.05 as significant (92). Genes with multiple barcodes were removed from downstream analysis. In total, barcodes representing 4,508 genes were analyzed across the two batches. Fitness scores correspond to the log_2_(fold change) output from edgeR. Hierarchical clustering was performed using Cluster 3.0 (93) and visualized using Java TreeView (94). Batch 1 and Batch 2 experiments were analyzed separately, comparing aneuploid fitness scores to batch-paired euploids unless otherwise noted. 5 generation collections (Chr4 *ssd1Δ*, Chr4 wild-type, and isogenic euploids) were analyzed separately. To maximize available statistical power, we combined and reanalyzed all seven euploid replicates from Batch 1 and Batch 2.

#### Enrichments

Enrichments for GO terms were assessed using setRank version 1.0 (95) or GO Term Finder version 0.86 in the S*accharomyces* Genome Database (96), compared to a background dataset of all measured genes. Statistical significance was assessed using Mann-Whitney U tests for continuous data and hypergeometric tests for categorical terms. DNA motif searches were performed on sequences -500 bp upstream and +100 bp downstream each gene’s open reading frame, using MEME suite (97) for novel motif discovery and SEA (98) to look for enrichment of previously identified motifs (22, 24). Propensity for disorder was quantified by % residues with IUPRED3 disorder propensity score >0.5 (99). Ssd1 targets were compiled from (21, 24). Other mRNA/protein features analyzed included those listed in (32) and age-dependent ribosome pause sites (100, 101).

#### Modeling

Linear modeling was performed using ordinary least squares (OLS) regression as implemented in the statsmodels version 0.11.1 package for python3. Visual representations of the regressions were created using seaborn version 0.11.2, with 95% confidence interval plotted using the lmplot function.

#### Relatively detrimental/beneficial gene sets

Unless otherwise noted, detrimental/beneficial gene sets correspond to barcodes with statistically significant decreases/increases in abundance (log_2_(fold change) < 0 or > 0, respectively, and FDR < 0.05). The 218 genes described in the text were identified comparing 7 replicates of wild-type and *ssd1Δ* euploid cells (FDR < 0.05). Genes more detrimental in aneuploids than euploid were taken as those significantly detrimental in aneuploids (FDR < 0.05) and >1.5 fold more deleterious in each aneuploid. To identify especially beneficial genes in *ssd1Δ* aneuploids, we required a significantly beneficial effect (logFC > 0, FDR < 0.05) and a fitness score difference of >= 0.5 in at least one *ssd1Δ* strain as compared to euploid wild-type, euploid *ssd1Δ*, and the corresponding wild-type aneuploid, if available. These criteria were used to generate the starting list of 445 especially beneficial genes. We relaxed criteria to identify overlapping genes shown in Fig 5B if genes in this set displayed a significantly positive log_2_ fitness score in other *ssd1Δ* aneuploid strains.

## Acknowledgements

Thanks to Michael Place for technical assistance and to the Gasch Lab for helpful comments. This work was funded by NIH grants R01CA229532 and R01GM148975 to APG. HAD was supported by training grants T32GM007133 and T32HG002760 to the Genomic Sciences Training Program.

## Author Contributions

H.A.D and A.P.G. designed research; H.A.D., J.H. H.H., J.R. performed research; H.A.D. analyzed data; H.A.D. and A.P.G. wrote the paper.

## SUPPLEMENTAL FIGURES

**Suppl. Fig 1.**
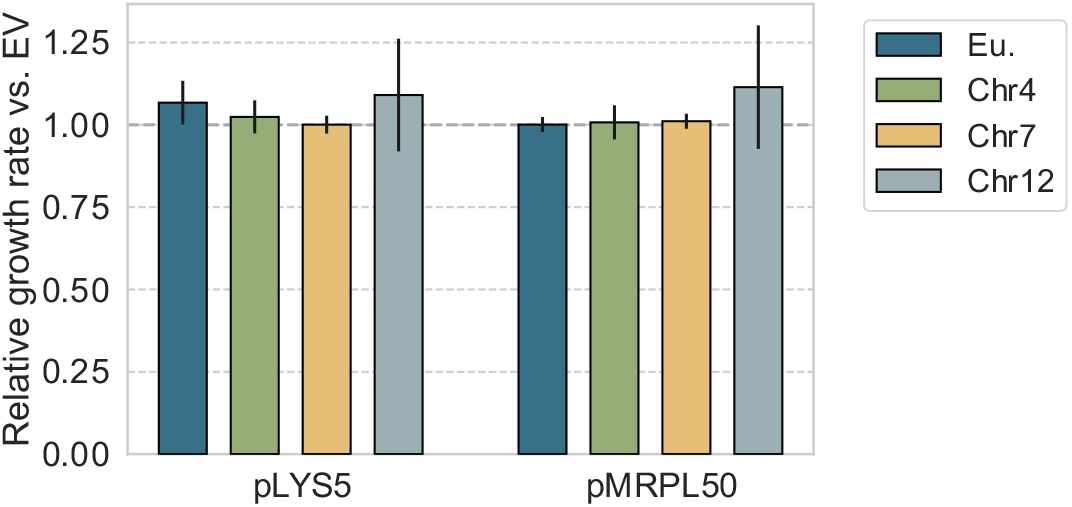
Genes scored as neutral in the barcoded library are validated as neutral when strains are grown in isolation. Two genes that are confidently scored as neutral in the library experiment were selected for investigation. The figure shows the average and standard deviation of relative growth rates of each strain harboring the indicated MoBY 1.0 plasmid vs. empty vector (EV), grown in selective media (SC-His + NTC + G418), n >= 3. The experiment confirms that these plasmids are indeed neutral and validate the library normalization procedure applied in this study.

**Suppl. Fig 2.**
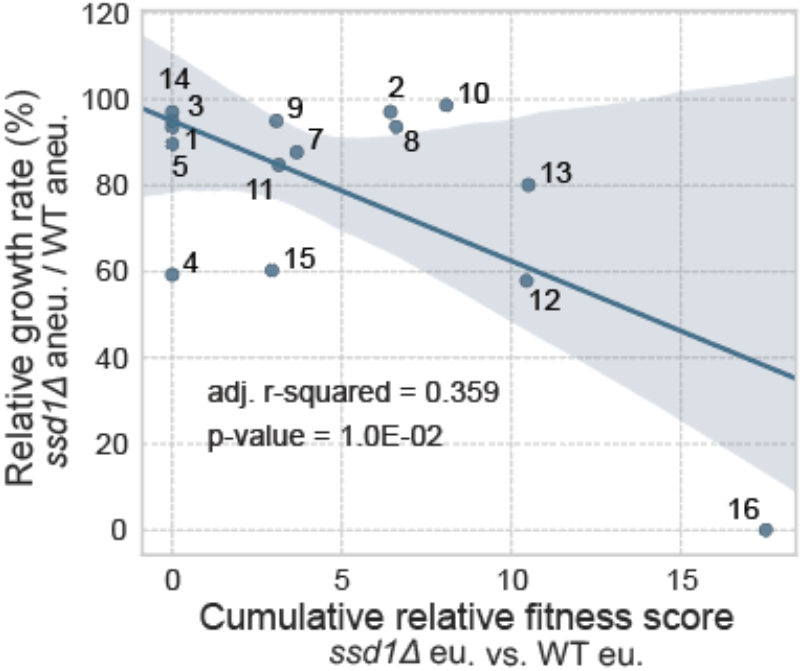
Linear modeling using fitness for a small handful of genes. Average relative growth rates of each *ssd1Δ* aneuploid versus corresponding wild-type aneuploid (y-axis, as shown in Fig 1) plotted against the sum of replicate-averaged relative fitness scores (*ssd1Δ* euploid versus SSD1+ euploid) for the 10% most relatively toxic genes in *ssd1Δ* euploid relative to wild-type euploid (x-axis).

**Suppl. Fig 3.**
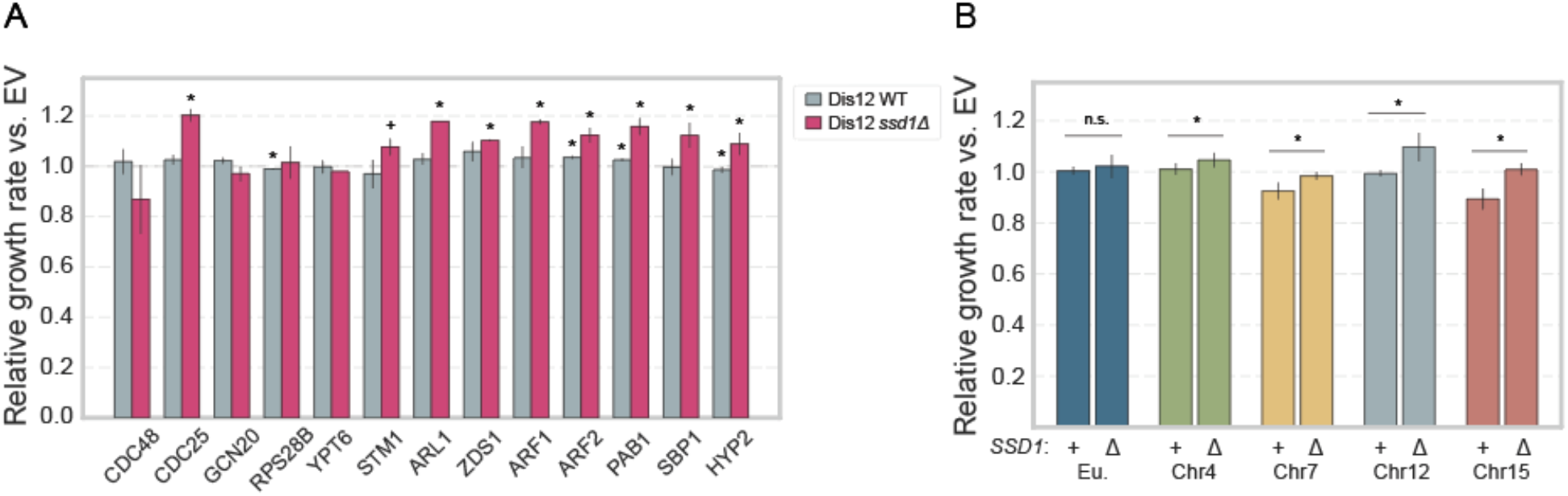
Duplication of select plasmids relatively improves growth rate of *ssd1Δ* aneuploids compared to *SSD1+* aneuploids. A) Average and standard deviation of growth rates for the denoted MoBY 1.0 gene duplication plasmid vs. empty vector (EV) in Chr12 aneuploids (n>=2) grown in rich media (YPD + G418). (*) indicates p < 0.0.05 of one-tailed, paired t-test between denoted strain and EV; (+) indicates p < 0.1. B) Average and standard deviation of growth rates for the HYP2 plasmid vs. empty vector in the indicated strains in rich media (n =3); (*) indicates p < 0.05 from one-tailed, paired t-test comparing relative growth rates between *SSD1+* and *ssd1Δ* strains.

